# Toward a unified connectomic target for deep brain stimulation in obsessive-compulsive disorder

**DOI:** 10.1101/608786

**Authors:** Ningfei Li, Juan Carlos Baldermann, Astrid Kibleur, Svenja Treu, Harith Akram, Gavin J.B. Elias, Alexandre Boutet, Andres M. Lozano, Bassam Al-Fatly, Bryan Strange, Juan Barcia, Ludvic Zrinzo, Eileen Joyce, Stephan Chabardes, Veerle Visser-Vandewalle, Mircea Polosan, Jens Kuhn, Andrea A. Kühn, Andreas Horn

**Author notes:** **Corresponding Author**, Ningfei Li, Department of Neurology, Movement Disorders and Neuromodulation Unit, Charité - University Medicine (CCM), Berlin, Germany.

## Abstract

Multiple surgical targets have been proposed for treating obsessive-compulsive disorder (OCD) with deep brain stimulation (DBS). However, different targets may modulate the same neural network responsible for clinical improvement. Here we analyzed data from four cohorts of OCD patients (N = 50) that underwent DBS to the anterior limb of the internal capsule (ALIC), the nucleus accumbens (NAcc) or the subthalamic nucleus (STN). Fiber tracts that were predominantly connected to electrodes in good or poor DBS responders were isolated from a normative structural connectome and assigned a predictive value. Strikingly, the same fiber bundle was related to treatment response when independently analyzing two large training cohorts that targeted either ALIC or STN. This discriminative tract is a subsection of the ALIC and connects frontal regions (such as the dorsal anterior cingulate, dACC, and ventral prefrontal, vlPFC, cortices to the STN). When informing the tract solely based on one cohort (e.g. ALIC), clinical improvements in the other (e.g. STN) could be significantly predicted, and vice versa. Finally, clinical improvements of eight patients from a third center with electrodes in the NAcc and six patients from a fourth center in which electrodes had been implanted in both STN and ALIC were significantly predicted based on this novel tract-based DBS target. Results suggest a functional role of a limbic hyperdirect pathway that projects from dACC and vlPFC to anteriomedial STN. Obsessive-compulsive symptoms seem to be tractable by modulating the specific bundle isolated here. Our results show that connectivity-derived improvement models can inform clinical improvement across DBS targets, surgeons and centers. The identified tract is now three-dimensionally defined in stereotactic standard space and will be made openly available.

---

Obsessive-compulsive disorder is a debilitating disease with a life-time prevalence of around 2.3%^1^. Treatment of severe cases by deep brain stimulation (DBS) to the ALIC has been approved by the FDA (Humanitarian Device Exemption) in 2009^2^. A variety of other targets have been proposed, however, including the STN^3,4^, the NAcc^5–7^, ventral capsule/ventral striatum (VC/VS)^8^, inferior thalamic peduncle (ITP)^9,10^, bed nucleus of the stria terminalis (BNST)^11^, anteromedial globus pallidus interna (amGPi)^12^, superolateral branch of the medial forebrain bundle (slMFB)^13^ and medial dorsal and ventral anterior nuclei of the thalamus (MD/vANT)^14^ (for an overview of targets see^15^). A recent prospective clinical trial implanted four electrodes per patient with one pair in the STN and one in the ALIC^16^.

In parallel, DBS has experienced a conceptual paradigm-shift away from focal stimulation of specific brain nuclei (such as the subthalamic nucleus or globus pallidus in Parkinson’s Disease; PD) toward modulating distributed brain networks (such as the motor basal-ganglia cortical cerebellar loop in PD)^13,17–23^. While this concept of modulating white-matter tracts (instead of grey matter nuclei) is certainly not new (and anterior capsulotomy was introduced already in the ~1950ies by Talairach and Leksell^24^), novel MRI technologies such as diffusion-weighted imaging based tractography have been used in functional neurosurgery in order to more deliberately target white-matter tracts^21^. In this translational development, the Coenen and Mayberg groups should be explicitly mentioned, among others, for pioneering and rapidly translating the use of tractography to functional surgery since around 2009^13,18–20,25,26^.

Thus, it could be possible that, of the multiple targets proposed, some – or most – may in fact modulate the same brain network to alleviate symptoms. Such a concept has been proposed in the past by Schlaepfer and colleagues for the case of treatment-refractory depression^27^. According to their concept, the supero-lateral branch of the medial forebrain bundle may connect most if not all surgical targets that were proposed for treatment of depression (e.g. subgenual cortex, ALIC, NAcc, habenula). Thus, in theory, the *tract itself* could be a surgical target – and could be modulated in a similar way when targeting various points along its anatomical course. Accordingly, already, Coenen and colleagues surgically implanted electrodes in two OCD patients, targeting a tract instead of a localized target^13^. The tract connected the ventral tegmental area and the prefrontal cortex and authors referred to it as the superolateral branch of the medial forebrain bundle.

Other invasive therapies, such as cingulotomy and capsulotomy also aimed at disrupting connectivity from frontal regions by lesioning white matter bundles^28^. It could recently be shown that such tract- or network-based concepts may be used to predict clinical improvements across DBS centers and surgeons for the case of Parkinson’s Disease^29^. Based on modern neuroimaging methods and high-resolution connectomic datasets, connectivity of DBS electrodes to specific cortical regions was associated with stronger therapeutic effects in various diseases treated with this surgical procedure^29–33^.

For the case of OCD, Baldermann and colleagues recently demonstrated that structural connectivity from DBS electrodes to medial and lateral prefrontal cortices were associated with stronger symptom alleviation^30^. Crucially, they were also able to identify a specific subsection of the ALIC that was highly associated with symptom improvements after one year of DBS. Of note, connectivity to this fiber tract was able to predict ~40 % of the variance in clinical outcome in out-of-sample data. The bundle was described to connect to both the medial dorsal nucleus of the thalamus and to the anterior part of the STN (which both have received substantial attention in the context of OCD). The STN itself is a prominent target for DBS of various diseases including PD^34^, dystonia^35^, OCD^36^ and Tourette’s Syndrome^37^. The small nucleus receives wide-spread direct afferents from most parts of the prefrontal cortex and is involved in motor, associative and limbic processing^38^. Due to these spatially organized cortico-subthalamic projections, the nucleus has various functional zones that largely follow the organization of the frontal cortex, i.e. the sensorimotor parts of the STN are situated posterior and followed by pre-/oculomotor-, associative and limbic domains in anteromedial direction.

Consequently, the anterior (associative/limbic) parts of the STN have been targeted by DBS for OCD^36^; these same anterior subregions were exclusively connected to the tract-target identified by Baldermann et al. in ALIC-DBS patients^30^. Following up on this, our present study aimed at testing whether the same tract could be associated with good clinical outcome in a cohort treated with STN-DBS. We retrospectively analyzed two cohorts of DBS patients that were treated with either STN-DBS or ALIC-DBS in order to test our hypothesis, that the same tract could potentially predict clinical improvement in STN-DBS as well as ALIC-DBS. In this attempt, we identified a common tract that already became apparent when analyzing either cohort alone. After calculating the tract exclusively based on data of one cohort (e.g. ALIC), we cross-predicted outcome in the other cohort (e.g. STN), and vice versa. We then tested predictive utility of this tract in two additional cohorts from a third and fourth center. Finally, we set the resulting tract target into the larger context of OCD-DBS literature and tested, whether it could be used to explain outcomes of reported clinical studies with different surgical targets.

## Results

Two cohorts (Cologne; ALIC target; N = 22; and Grenoble; STN target; N = 14, two electrodes in each patient) formed a training and cross-validation sample in which the tract target was identified and validated. Each of the two cohorts were first analyzed independently, then used to cross-predict outcome in patients from the other one. The main part of our analyses focuses on these two cohorts. As further validation of results, two additional test-cohorts were included (Madrid: two electrodes in each patient targeting Nucleus Accumbens (NAcc); London: four electrodes in each patient targeting both ALIC and STN).

Patients in all cohorts were of similar age with a similar Y-BOCS score at baseline and comparable Y-BOCS improvement scores (Table 1). In the first test cohort (Madrid; NAcc target; N = 8), improvement scores were taken after activating each of the four electrode contact pairs for 3 months, respectively (following the clinical protocol as described in^39^). This resulted in a total of 32 data points. In the second test cohort (London; both ALIC and STN target; N = 6, four electrodes in each patient), stimulation parameters resulted from an optimized phase following parameter optimization.

**Table 1:** Patient demographic details and clinical results of the two cohorts

| | ALIC DBS Cohort<br>(Mean ± SD) | STN DBS Cohort<br>(Mean ± SD) | NAcc DBS Cohort<br>(Mean ± SD) | Combined DBS Cohort<br>(Mean ± SD) |
| --- | --- | --- | --- | --- |
| Center | University Hospital Cologne | University Hospital Grenoble | Hospital Clínico San Carlos Madrid | University Hospital London |
| Reference(s) | (22, 31) | (28) | (40) | (16) |
| N patients (females) | 22 (12) | 14 (9) | 8 (4) | 6 (1) |
| N electrodes | 44 | 28 | 16 | 24 |
| Age | 41.7 ± 20.5 | 41 ± 9 | 35.3 ± 10.4 | 45.5 ± 10.5 |
| Y-BOCS Baseline | 31.3 ± 4.4 | 33.4 ± 3.7 | 30 ± 7.75 | 36.2 ± 1.8 |
| Y-BOCS after DBS | 20.7 ± 7.7<br>(12 months postop) | 19.6 ± 10.6<br>(12 months postop) | 14.75 ± 7.2<br>(3 months postop of best contact) | 14.3 ± 4.1<br>(optimized phase in [16]) |
| Absolute Y-BOCS Improvement | 9.6 ± 6.5 | 13.8 ± 10.8 | 15.1 ± 9.6 | 21.83 ± 5.7 |
| % Y-BOCS Improvement | 31.0 ± 20.5 % | 41.2 ± 31.7 % | 47.8 ± 23 | 50.0 ± 12.6 % |

Electrode localization confirmed accurate placement to each of the three target regions for all patients of the four cohorts (Figure 1).

**Figure 1.**
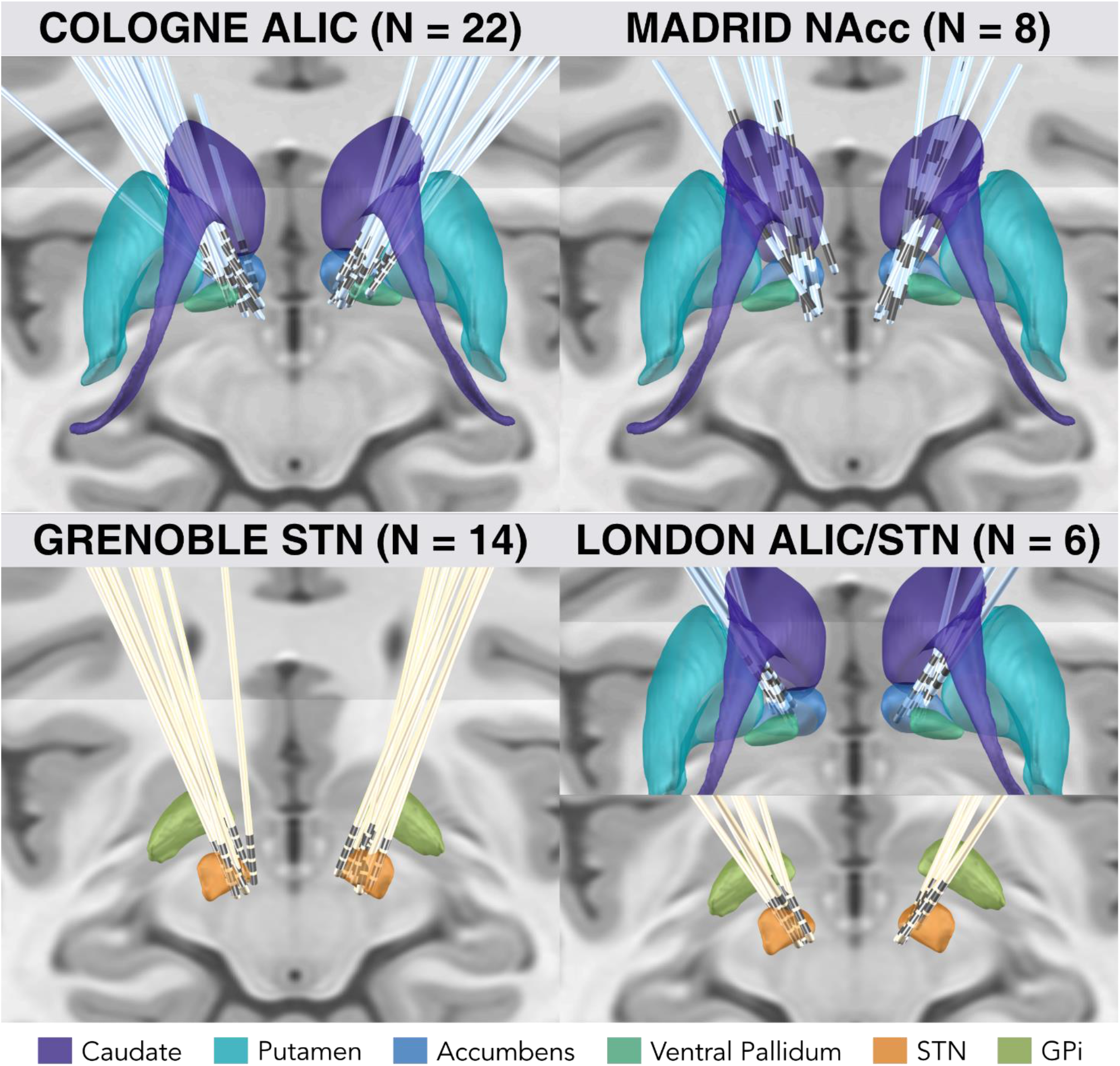
Overview of lead electrodes placement of the two training/cross-validation cohorts (left) targeting ALIC DBS (Cologne) and STN DBS (Grenoble) cohorts, and the two test cohorts (right) targeting NAcc (Madrid) and both ALIC & STN with four electrodes per patient (London). Subcortical structures defined by CIT168 Reinforcement Learning Atlas^40^ (ALIC/NAcc region) and DISTAL Atlas^41^ (STN region), with coronal and axial planes of the T1-weighted ICMB 152 2009b nonlinear template^42^ as background.

Connectivity analysis results seeding from electrodes of the two training cohorts (Cologne and Grenoble) based on the N = 985 HCP normative connectome are shown in Figure 2. The overall connectivity of electrodes to other areas in the brain (without weighing for clinical improvement) was strikingly different between the two cohorts (Figure 2, top row). This is hardly surprising since it mainly reflects the overall structural connectivity profiles of the two DBS targets and the STN as a widely connected basal ganglia entry point and the ALIC as a white matter structure are differently connected in the brain. However, when tracts were weighted by their ability to discriminate between good and poor responders (using the Fiber T-score method described below), a bilateral positively discriminative tract to the medial prefrontal cortex emerged in each cohort even when cohorts were analyzed independently (Figure 2, middle row). The degree of lead connectivity to this tract correlated with clinical improvement (R = 0.63 at p < 0.001 in the ALIC cohort and R = 0.77 at p < 0.001 in the STN cohort; Figure 2, bottom row).

**Figure 2.**
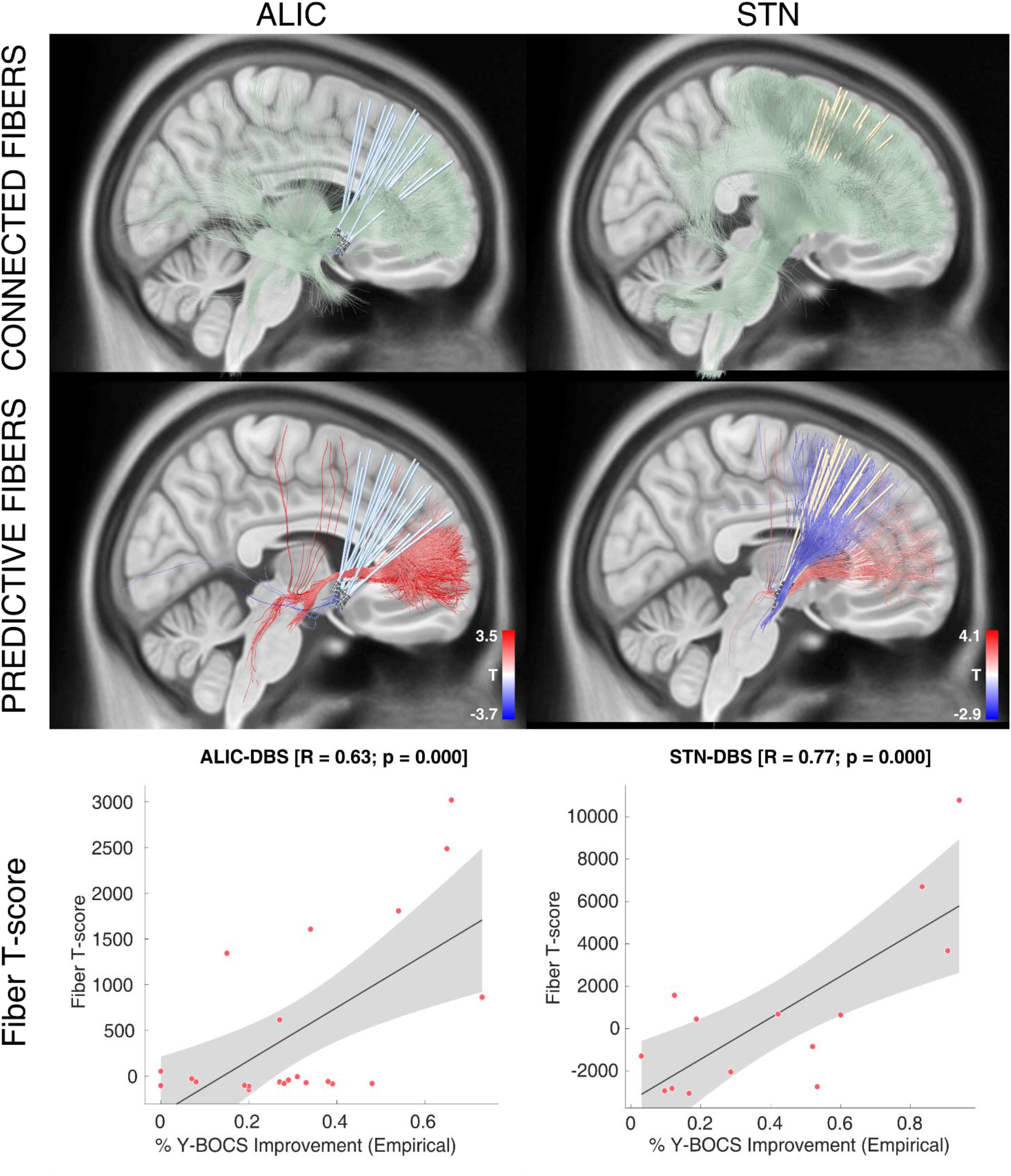
Predictive fiber tracts in training cohorts. Top: All fibers connected to the sum of VTAs of each cohort are shown in green. Middle: Predictive fibers positively associated with clinical improvement are shown in red, while those negatively associated with clinical improvement are shown in blue. The top and bottom 20% (based on Fiber T-scores) of the predictive fibers are displayed. Bottom: Correlations between the degree of stimulating positively predictive tracts (sum of aggregated Fiber T-scores under each VTA) and clinical improvements. While this analysis is based on a normative connectome, a replication of it based on anatomically predefined pathways is shown in figure S1.

Of note, these correlations are somewhat circular and meant to describe the degree of how well discriminative tracts could explain the same sample of patients on which they had been built. More interestingly, in the next step, the tract was calculated exclusively on data from the STN cohort and then used to explain outcome in the ALIC cohort (R = 0.50 at p = 0.009) and vice versa (R = 0.49 at p = 0.041) (Figure 3).

**Figure 3.**
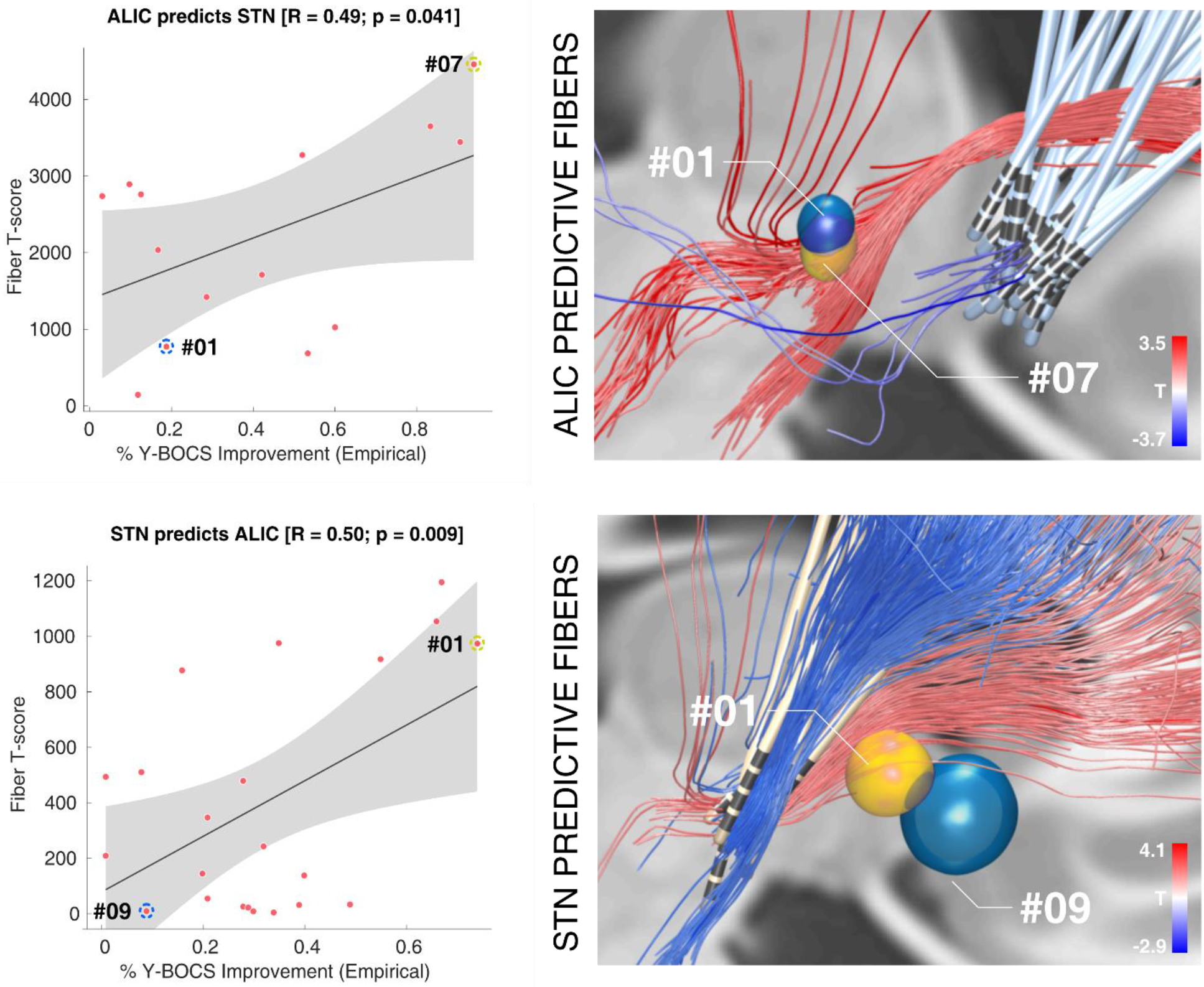
Cross-prediction between the ALIC and STN training cohorts. Top: When the tract was calculated exclusively based on data from the ALIC cohort, it was used to calculate Fiber T-scores for all patients in the STN cohort. These were correlated with clinical improvements in the STN cohort. One example patient with strong overlap of the tract (yellow) received a high Fiber T-score, whereas one with less overlap received a lower score (blue). The two example patients are marked in the correlation plot on the left. Bottom: Here, the tract was calculated exclusively on data from the STN cohort to predict outcome in patients in the ALIC cohort. Again, two example patients are shown. Of note, here, some VTAs barely overlapped with the tract and consequently received a near-zero score.

Crucially, some VTAs of the ALIC cohort resided entirely below the identified tract and thus received a Fiber T-score of (near) zero (also see blue example patient in Figure 3, bottom right). The same holds true when either calculating the tract based on the STN cohort (Figure 3) or the ALIC cohort itself (Figure 2). To further investigate this matter, two-sample t-tests between improvements of patients with near zero scores (Fiber T-scores below 50) and the remaining patients with VTAs covering the tract well (scores above 50) were calculated. This showed that electrodes that reached the tract well resulted in significantly better clinical improvement (T = 6.0 at p < 10^−5^ when the tract was calculated on the ALIC cohort, Figure 2, and T = 3.7 at p < 0.005 when it was calculated on the STN cohort, Figure 3).

Depending on the target, the analysis revealed different proportions of “positive” and “negative” fibers (ALIC cohort: 22.2k positive vs. 1.9k negative fiber tracts selected from the group connectome; STN cohort: 45.1k positive vs. 48.6k negative fibers and both cohorts combined: 54.4k positive vs. 9.6k negative fibers).

In the next step, the analysis was performed on the two cohorts combined. Again, the same tract emerged, now even more clearly (Figure 4, top). Thus, bundles were highlighted, that were predominantly connected with VTAs of patients from both cohorts with good or poor improvement, respectively. The resulting positive discriminative tract traversed slightly dorsal to the group of electrodes of the ALIC-cohort and coursed centrally or slightly ventral to the electrodes of the STN cohort. This tract was then used to predict outcome in two completely independent test-cohorts of patients that underwent surgery in a third and fourth center (Madrid & London; Figure 4, bottom). While the surgical target of the Madrid cohort was the Nucleus Accumbens, electrode placement is comparable to the ALIC / Cologne cohort (Figure 1). Here, improvements were taken for each contact pair during a three-month interval, leading to 32 data points (Figure 4, bottom left, active contact pair color coded). In the London cohort, patients had received two electrodes to each target (four in total) and scores were summed up across targets. In both test-cohorts, stimulation overlap with the tract target significantly correlated with empirical improvement (Madrid: R = 0.50 at p < 0.001, London: R = 0.75 at p = 0.040). Of note, VTAs in the London sample were estimated with a different software (see methods), patients received four electrodes and the clinical scores represented an “optimized” phase following nine months of a clinical trial^16^.

**Figure 4.**
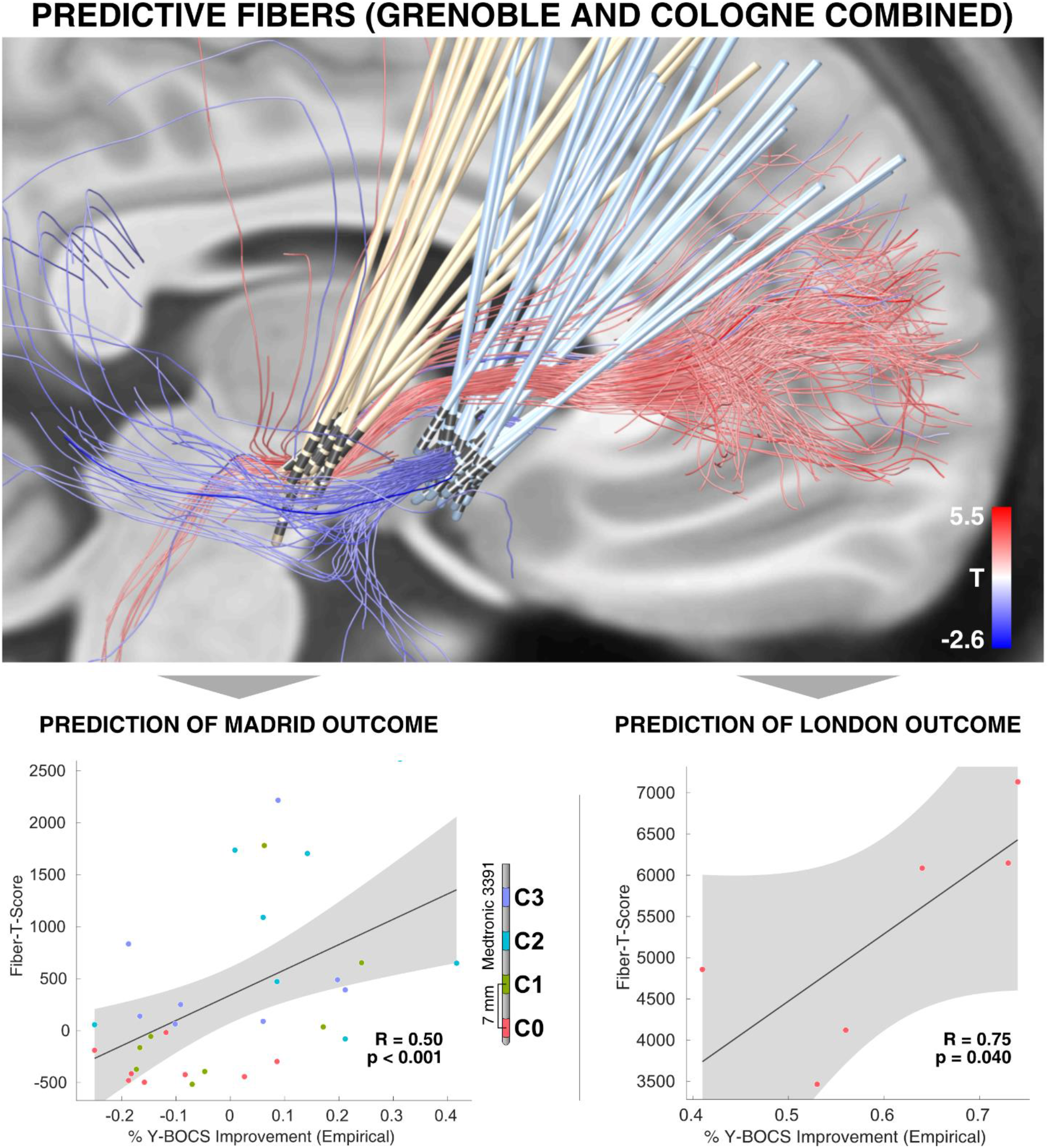
Predictions in test-cohorts. Top: predictive fibers calculated on both training cohorts (Cologne & Grenoble) irrespective of their target. Red fibers are positively associated with clinical improvement, blue fibers negatively. Bottom: the sum of aggregated Fiber T-scores under each VTA predicted %-Y-BOCS improvements in eight patients with 4 settings each (N = 32 stimulations) of the Madrid cohort (left) and six patients of the London cohort with dual stimulation of STN and ALIC (right). Please note that the p-values in this manuscript are based on random permutation testing. Based on classical tests, the result shown in the lower right panel would remain significant in a one-sided test, only (p-one-sided = 0.044, p-two-sided = 0.089). A replication of this result based on anatomically predefined pathways may be found in Figure S2.

Given the high amount of false-positive connections that are present in dMRI based connectomes^43^, we replicated all findings of the study using a synthesized anatomical atlas that is purely based on established anatomical knowledge^22^ and thus free of such falsepositive connections. Results that were based on this atlas were highly similar and isolated the hyperdirect pathway connecting the dorsal anterior cingulate cortex (dACC) to the STN to be positively predictive of clinical outcome (Figures S1–2).

The tract target identified here may potentially “unify” some aspects of the STN and ALIC/NAcc targets for OCD. Thus, in a final analysis, we aimed at setting it into context with other DBS targets that were used in OCD-DBS, before. To do so, we converted literature-based targets into MNI space^44^ and set them into relation with the tract target (see Figure 5, Table 2 and supplementary material). A large number of reported DBS targets for OCD seemed to cluster on or around it. Furthermore, clinical improvement values that had been reported in these studies could be significantly explained by calculating the weighted overlap between stereotactic target sites and the tract (Figure 5 C, see supplementary material for details).

**Figure 5.**
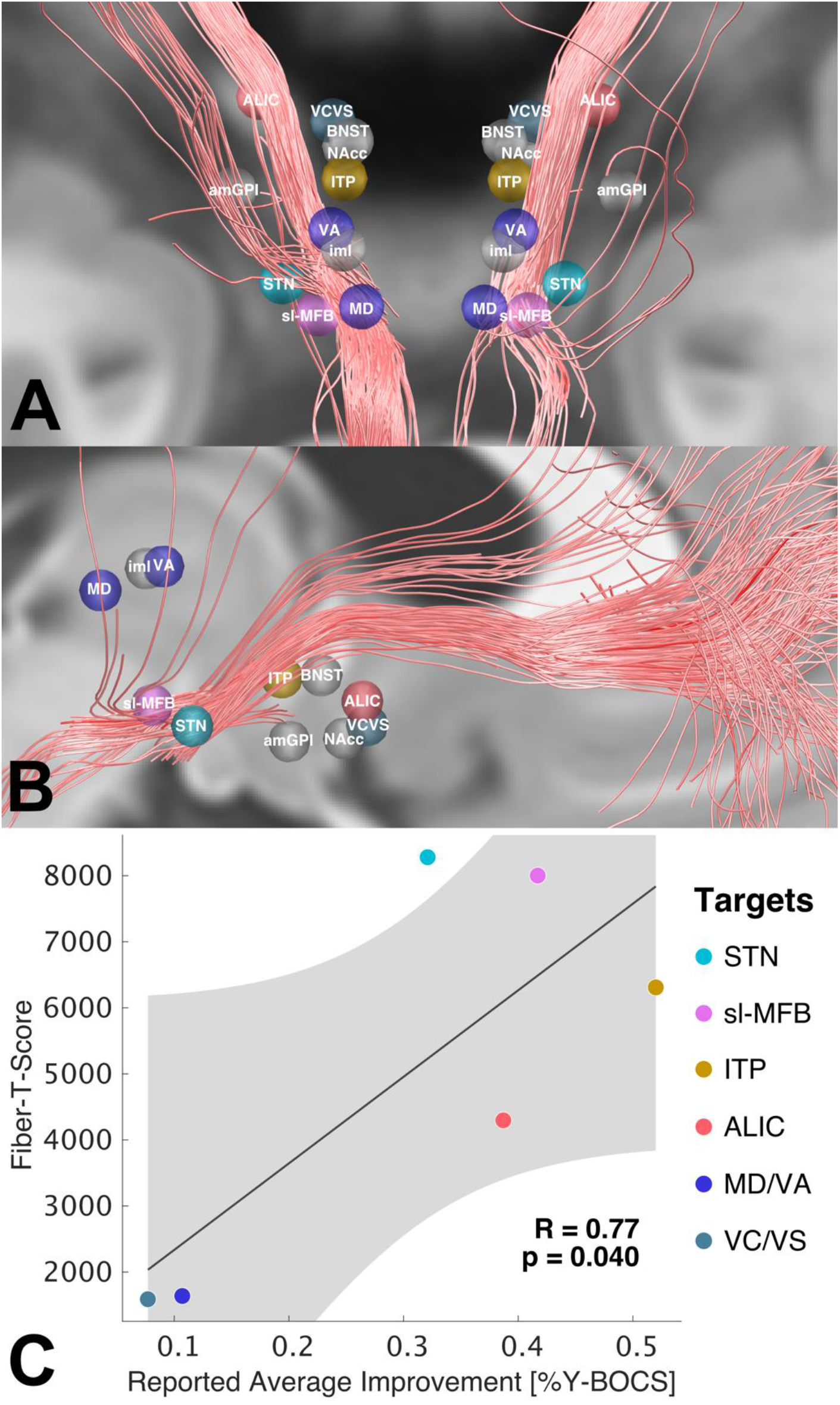
Overview of the positively predictive fiber tracts identified in the present study in synopsis with DBS targets for treatment of OCD based on literature results. Note that most targets were reported for the tip of the electrode, thus, active stimulation may have occurred dorsal to shown targets (Table 2). A & B show reported average stimulation sites mapped to standard space. C shows the degree of weighted overlap between stimulation sites and the identified tract. These were correlated with reported average %-Y-BOCS improvements of published studies (where available, other sites marked in gray; see supplementary material for details).

**Table 2:** DBS targets for treatment of OCD defined in the literature. *MD*, medial dorsal thalamic nucleus; *VA*, ventral anterior thalamic nucleus; *iml*, internal medullary lamina; *MCP*, mid-commissural point; *AC*, anterior commissure * Tourette patients, with prominent symptoms of OCD

| DBS Target | Reference | Number of patients | % YBOCS change | AC/PC coordinates | Relative to | Target type | MNI coordinates (Figure 5) |
| --- | --- | --- | --- | --- | --- | --- | --- |
| STN | Mallet et al. 2008<sup>50</sup> | 8 | 32.1 | NA | AC | Tip of the electrode | ±11.30 -9.90 -7.81 |
| amGPi | Nair et al. 2014<sup>12</sup> | 4* | NA | ±14.47 9.85 -3.28 | MCP | Tip of the electrode | ±15.66 -1.41 -8.22 |
| VC/VS | Tsai et al. 2010<sup>51</sup> | 1 | 7.7 | ±7.5 16.3 -3.05 | MCP | Tip of the electrode | ±7.92 5.51 -9.01 |
| sl-MFB | Coenen et al. 2017<sup>13</sup> | 2 | 41.7 (at 12 months) | ±7.6 -1.72 -3.0 | MCP | Active contacts | ±8.35 -13.64 -7.00 |
| NAcc | Sturm et al. 2003<sup>5</sup> | 4 | NA | ±6.5 2.5 -4.5 | AC | Tip of the electrode | ±6.98 3.69 -10.55 |
| ALIC | Nuttin et al. 2003<sup>52</sup> | 6 | 38.7 | ±13 3.5 0 | AC | Tip of the electrode | ±13.84 5.17 -5.04 |
| MD | Maarouf et al. 2016<sup>14</sup> | 4 | 10.7 | ±4.7 -18.52 4.87 | AC | Active contacts | ±5.10 -18.17 2.59 |
| VA |  |  |  | ±6.84 -13.76 7.78 |  |  | ±7.52 -12.68 5.60 |
| iml |  |  |  | ±5.78 -14.9 7.08 |  |  | ±6.36 -13.99 4.85 |
| ITP | Lee et al. 2019<sup>10</sup> | 5 | 52.0 | ±6.5 -3 -0.5 | AC | Tip of the electrode | ±6.92 -1.84 -5.13 |
| BNST | Nuttin et al. 2013<sup>53</sup> | 4 | NA | ±6 0 0 | AC | Tip of the electrode | ±6.33 1.39 -4.87 |

Given the potential clinical importance of the identified tract, we characterized its anatomical properties using additional views relative to anatomical landmarks (Figures 6 & S3) as well as in comparison to anatomical dissection results (Figure S4). Anatomically, the tract is a subpart of the well-characterized ALIC that connects areas of the prefrontal cortex with the subthalamic nucleus and MD nucleus of the thalamus^45,46^. Anatomical validity of the isolated tract was discussed with four anatomists (see acknowledgement section). In the motor domain, the “hyperdirect pathway”, i.e., a direct connection from the frontal cortex to the subthalamic nucleus, has been well established^47,48^, functionally, but the STN is known to receive widespread and direct input from most areas of the prefrontal cortex^45^. Thus, the main part of the specific bundle delineated here may represent a route of direct input from frontal regions to the STN. In addition, connections between mediodorsal nucleus of the thalamus and prefrontal regions received slightly lower T-scores and are not shown in 3D visualizations but well visible in 2D sections shown in Figure 6.

**Figure 6.**
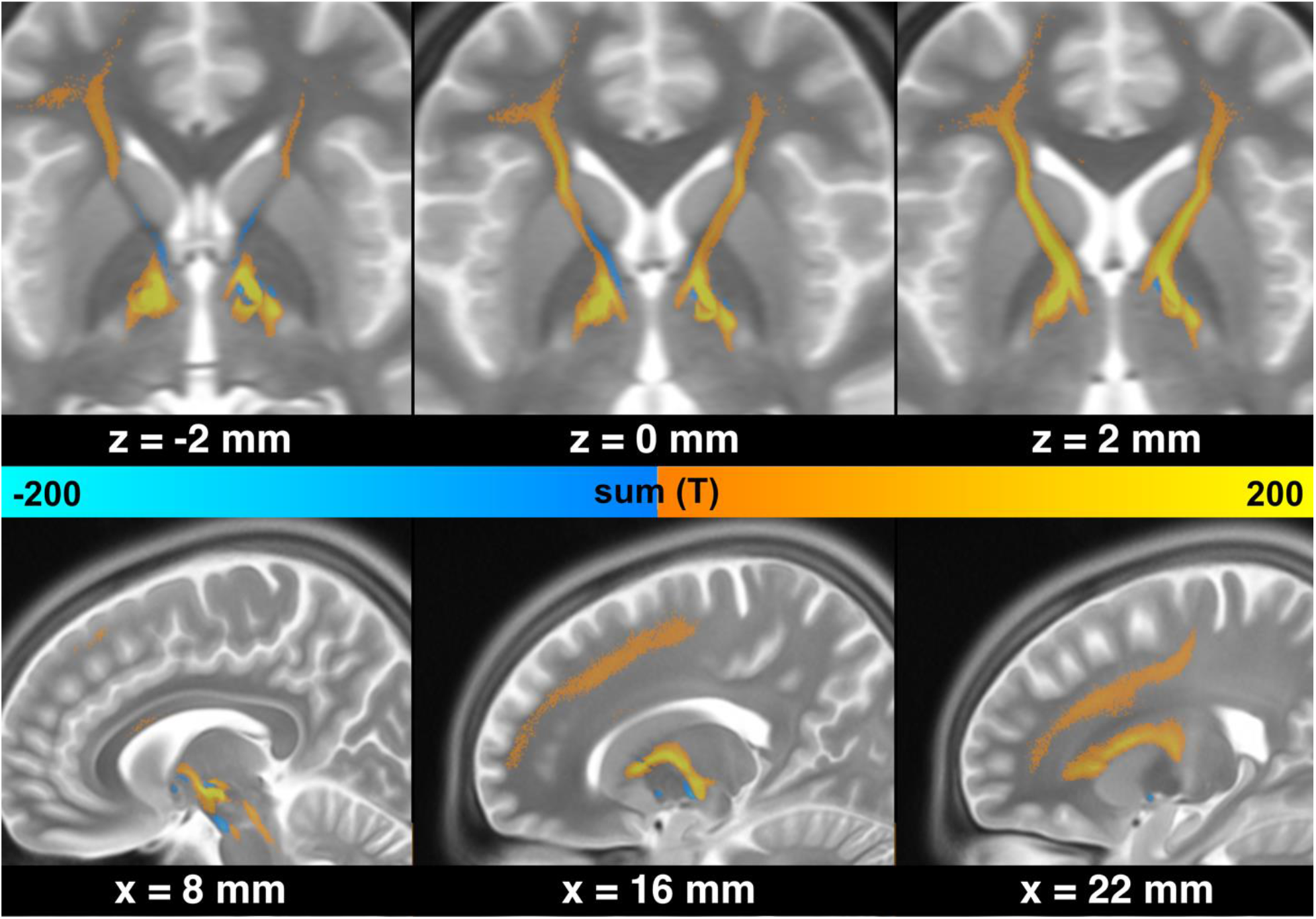
Anatomical course of peaks in discriminative fibers shown in MNI space. The tract is connected to the subthalamic nucleus and mediodorsal nucleus of the thalamus, traverses through the anterior limb of the internal capsule and has a wide array of frontal connections including dorsal anterior cingulate cortex and ventrolateral prefrontal cortex.

To properly define the anatomical course of this tract, we openly released it as an atlas in stereotactic (MNI) space within Lead-DBS software (www.lead-dbs.org). Of note, Lead-DBS is scientific and not clinical software and the tract should not be vacuously used for any form of clinical decision making^49^.

## Discussion

We analyzed data from four cohorts of OCD patients with different DBS targets using a connectomic approach. Strikingly, the same optimal tract target emerged when separately analyzing either of a large ALIC-DBS and STN-DBS cohort, alone. Among other connections, this bundle connected the dorsal anterior cingulate and ventrolateral prefrontal cortices to the anteriomedial STN. When the tract was calculated on each cohort alone, it could be used to cross-predict clinical improvement in the other cohort. Furthermore, clinical outcomes in two independent test cohorts from a third and fourth center could be significantly predicted based on overlap with the tract. Finally, literature-based stimulation sites for OCD seemed to cluster close to the identified tract. Indeed, their spatial overlap with the tract correlated with reported clinical improvement across studies.

In recent work by Baldermann et al. 2019^30^, a fiber bundle was identified that was associated with good clinical improvement in ALIC-DBS for OCD when modulated by high-frequency DBS. This bundle connected the ventrolateral prefrontal cortex, dorsal anterior cingulate cortex, subthalamic nucleus and mediodorsal nucleus of the thalamus. The authors concluded that DBS would modulate the communication between these sites, which is in line with the assumption that a dysregulation in the fronto-striato-thalamic circuit may pathophysiologically underlie OCD^54–56^.

Here, we extend anatomical definition of the same circuit and show that it emerges based on data from multiple stimulation sites. The subthalamic nucleus receives afferents from a large portion of the prefrontal cortex by hyperdirect pathways that are known to traverse within the internal capsule^45,57^. Recently, such an input to the STN from prefrontal regions was electrophysiologically described in humans^58^. In rodents, lesions to such a “limbic hyperdirect pathway” led to diminished discriminative accuracy and increased perseveration^59^. One classical cortical region which was described as an origin of limbic hyperdirect input is the dACC^22,45,60^ which crucially plays a prominent role in the classical cortico-striato-thalamo-cortical (CSTC) model of OCD^60^ and leads to improvement of OCD symptoms when directly lesioned in humans^61^. In a novel theory of cognitive control dysfunction, McGovern & Sheth attributed a central role to the dACC^60^. The normative connectome analysis identified the dACC as a cortical connection site to the identified tract, among others. Because of the high amount of false-positive connections in diffusion MRI based connectomes^43,62^, we repeated the analysis using an atlas of predefined anatomical tracts^22^. Here, the hyperdirect pathway connecting dACC to the STN was isolated as the only of five bundles in the ALIC that were included in the atlas (Figures S1–2). Thus, hyperdirect cortical input from dACC to STN could be an anatomical and functional substrate of the identified bundle. In this context, it is crucial to note that the atlas by nature cannot represent each and every white-matter bundle that exists in the ALIC / STN region and shows “gaps” in between the included bundles (Figures S1–2). Thus, while normative connectomes include a large number of false-positive fibers, the atlas may instead be prone to false-negative connections, since some tracts are simply not included. For instance, it is known that the STN receives direct input from other areas of the prefrontal cortex such as the ventrolateral prefrontal cortex^63^. In summary, while dACC and vlPFC are likely candidates to play a functional role, our methods and results are unable to determine the exact cortical region(s) of origin with absolute certainty. Despite this limitation, our results define a precise three-dimensional reconstruction of the tract itself (i.e. where exactly it traverses within the ALIC) in standard stereotactic space.

A highly similar pathway that already served as a tract-target in a small case-series of OCD patients^13^ also traversed within the ALIC but has instead been referred to as the superolateral branch of the medial forebrain bundle (sl-MFB)^49^. The original anatomical definition of the medial forebrain bundle (which was first defined in the rat) suggests a more ventral route connecting the ventral tegmental area to the olfactory cortex while bypassing the red nucleus medially^46^. In other words, the anatomical definition of the medial forebrain bundle does not traverse within the ALIC. This mismatch between the surgical target (sl-MFB) and anatomical literature (mfb) has recently been confirmed by the original authors of the surgical target and they now additionally referred to it with “vtaPP” (for ventral tegmental area projection pathway)^64^. This potentially misleading nomenclature of the surgical sl-MFB target has suggested that results in two previous OCD studies would be conflicting, while anatomically, their results agreed. Both studies favored a similarly defined tract within the ALIC, which was referred to as sl-MFB in one study^65^ and as anterior thalamic radiation in the second^30^. To readers, this suggested conflicting results while they were actually confirmatory (based on the location of both tracts within the ALIC). Thus, we welcome the recent steps taken to move away from calling the surgical target sl-MFB toward calling it “vta-PP”^64^. This said, our interpretation of the identified tract differs. Our findings reveal a tract connecting frontal areas with the STN (cf. Figure S3 C & results from the basal ganglia pathway atlas, Figure S1–2). Thus, we attribute the tract to afferents of the STN (limbic hyperdirect pathway) as opposed to efferents of the ventral tegmental area implied by the term “vtaPP”^64^.

This interpretation could be further supported by combined analyses of dMRI and tracing methods in nonhuman primates as well as human subjects, which were used to segregate prefrontal fibers passing through the internal capsule^66^. Fibers that originated from ventrolateral prefrontal cortices (areas 45 and 47) were shown to terminate in the medial part of the STN and the MD nucleus of the thalamus – precisely corresponding to the tract described here. Alternatively – or additionally – the hyperdirect pathway projecting from dACC to the STN may be functionally involved in mediating treatment outcome. As mentioned, a strong additional hint for this latter hypothesis is that lesions to the dACC itself have beneficiary effects on OCD^61^.

### Toward symptom-specific circuitopathies

Based on our results, two testable hypotheses with implications above and beyond OCD could be proposed. First, different surgical targets may reduce *the same* symptoms equally well – potentially by modulating the same tract or network. Second, in addition, they may modulate not only one (shared) network but other networks that are *not* shared, resulting in different changes across other behavioral domains. This can be seen by widely different connectivity profiles of the targets (Figure 2 top row) and differential effects of STN vs. ALIC stimulation on depressive / cognitive functions described by Tyagi et al.^16^. Thus, one may speculate that networks are symptom-specific (and not disease-specific). When modulated, these networks or tracts seem to not ameliorate a specific disease but rather specific symptoms present in the disease.

In OCD, accordingly, different symptom types (for example contamination vs. checking) were found to activate different prefrontal sites (ventromedial vs. dorsolateral, respectively)^67^. Similar observations have been made in other diseases, before. For instance, Akram and colleagues demonstrated that connectivity to specific cortical regions was associated with improvement in different clinical features of Parkinson’s Disease (e.g. connectivity to M1 preferentially reduced tremor while to the SMA reduced rigidity and bradykinesia)^31^. Similarly, connectivity from electrodes to M1 was associated with tremor improvement in Essential Tremor^68^.

Supporting the first hypothesis, our study was able to predict symptom-specific clinical improvement *across DBS targets and centers* based on connectivity data. While the tract that our data seems to shape out is predictive for Y-BOCS improvement, completely different tracts could have emerged when repeating the analyses for depressive or cognitive flexibility symptoms (as analyzed by Tyagi et al.). Unfortunately, these data were not available for the two main cohorts analyzed here.

Going further, shared symptom networks could be present in other diseases for which multiple surgical targets are investigated. Major Depression and Tourette’s Syndrome are obvious examples and extensive work in this direction is currently ongoing^19,69–72^. Similar concepts could even be applied to more established targets such as STN vs. GPi DBS^73–75^ or symptom-specific alleviations across diseases.

Potentially, DBS surgery in the (distant) future could involve detailed preoperative phenotyping to establish a broad patient-specific symptom score. Based on databases of clinical improvements along affected symptom axes, *a mix of networks* that would be modulated to alleviate each patient’s specific symptom profile could be identified. Such concepts are still mostly speculation but may be investigated by future studies. This said, we must emphasize that the present study investigated data on a group level and utilized connectivity from individuals without OCD. As mentioned by others in the very context, we could not agree more that surgical decision making for DBS should not be based on such aggregated normative data, alone^49^. Further studies are required to determine whether individual patient connectivity or generic connectome data (or both) could assist with optimizations in surgical targeting or DBS programming by determining crossing sites of symptom networks for specific patients.

### Limitations

Several limitations apply for the current work. First and foremost, the retrospective character of the study is not ideal to compare and study effects of clinical outcome which is why we kept clinical information to a minimum and instead referred to the underlying clinical studies. Second, we used normative connectome data instead of patient-specific diffusion-weighted MRI data (which is unavailable for most of the patients included). This poses dramatic limitations since such data cannot be representative of patient-specific anatomical variations. Still, we argue that some aspects about general pathophysiological mechanisms may still be investigated using normative data. Use of normative connectomes has been introduced in other clinical domains where patient-specific MRI data was unavailable, such as stroke^76–78^ or transcranial magnetic stimulation^79^. In DBS, the technique has been applied before and – as in the present study – has led to models that could be used to predict improvement in out-of-sample data^29,30,80^. In addition to the practical advantage of being applicable in cases where patient-specific data is lacking, normative data also has the theoretical advantage of much better data quality. In the present case, a connectome dataset was derived from a high N of 985 subjects scanned under research conditions by one of the best methodological groups in the world^81^. It may be logistically challenging to acquire data of such quality in a clinical routine setting (e.g. pre-operatively) in individual patients but will be feasible in specialized centers. Tractography based DBS targets pointed to coordinates that were sometimes >2 mm when repeating analyses on test-retest scans of the same subject^82^. However, patient-specific connectivity can never be reconstructed when using normative connectomes. Thus, normative connectomes will likely not embody the final solution to the connectomic surgery framework and will be challenged by advances in MRI technology and algorithm developments. Potentially, as a step in-between, using combined information from normative and patient-specific connectomes could embody a promising strategy that should be explored, in the future.

Inaccuracies in lead localization may result from the approach of warping electrodes into common space as done here. To minimize this issue, we used a modern neuroimaging pipeline that has been scientifically validated in numerous studies and involved advanced concepts such as brain shift correction^83^, multispectral normalization, subcortical refinement^83^ and phantom-validated electrode localizations^84^. The normalization strategy that was applied was found to automatically segment the STN as precisely as manual expert segmentations^85^ and each step of the pipeline was carefully assessed and corrected if needed by a team with long-standing expertise in this area^86,87^. Besides, both post-operative CT (thirty-three patients) and post-operative MRI (seven-teen patients) were used for electrode localization in the current dataset. Although studies have reported similar agreement between the results based on the two modalities, this might still lead to slight inconsistencies in the data. A larger dataset acquired with a homogeneous protocol would be ideal to validate our results in the future.

Importantly, given the correlative nature of the study, our findings may not differentiate between local and global effects. For instance, the tracts may have spuriously originated in the ALIC group because a more dorsal stimulation resulted with better clinical outcome. The congruency between results of the STN- and ALIC-cohorts resulting in the same fiber bundle still suggest that the identified tract could play a causal role. However, such a claim would need to be confirmed e.g. using optogenetics or electrophysiology.

It has been shown that dMRI-based tractography reconstructs a very high proportion of false-positive fibers in recent open challenges^43,62^. We aimed at reducing the risk of false positive tractography results in four ways. First, we used the tracking method that achieved the highest (92%) valid connection score among 96 methods submitted from 20 different research groups in a recent open competition^43^. Second, we used highest quality multi-shell diffusion data^81^ acquired on a high N (985 subjects) at a state-of-the-art imaging center (HCP data acquired at Washington University in St. Louis, see Acknowledgements). Third, we compared the tract results with anatomy text-books and discussed its validity with four anatomists (see Acknowledgements). Fourth, we replicated findings based on an atlas that is based on predefined anatomical tracts (see supplementary material). The tract described in the present study matches results from this atlas (Figure S1–2) and dissection studies (Figure S4). However, the potential that the tract represents a false positive result may not be completely ruled out given the fundamental limitations of dMRI-based tractography^43,62^.

## Conclusions

Four main conclusions may be drawn from the present study. First, we show that the overall connectivity profiles of STN- and ALIC-DBS electrodes project to largely different areas in the brain. Second, data in each target alone singled out the same fiber tract that was associated with long-term improvement of OCD symptoms when modulated either at the level of the STN or the ALIC. Third, we demonstrated that it is possible to cross-predict clinical improvement of OCD patients across DBS target sites (ALIC / STN) and centers (Cologne / Grenoble). Finally, we confirm results by predicting outcome in two additional cohorts from different centers (Madrid / London) and set results into context of published reports.

## Methods

### Patient Cohorts and Imaging

Fifty OCD patients from four centers were retrospectively enrolled in this study, among them twenty-two patients from University Hospital of Cologne implanted for ALIC DBS, fourteen patients from Grenoble University Hospital who underwent STN DBS surgery, eight patients who received bilateral electrodes targeting the NAcc from Hospital Clínico San Carlos in Madrid and six patients who received electrodes to both STN and ALIC from the National Hospital for Neurology and Neurosurgery. The patients from Cologne, Grenoble and Madrid received two electrodes each (N = 44 patients with N = 88 electrodes), the six patients in the London cohort received four electrodes each (N = 6 patients with N = 24 electrodes). All patients from Grenoble were bilaterally implanted with DBS electrodes 3389, as were all but three patients from Cologne, who received type 3387 electrodes (Medtronic, Minneapolis, Minnesota, US). Patients from London received models 3389 to the STN and 3387 to the ALIC. Patients from Madrid received models 3391. All patients qualified for DBS surgery based on their diagnoses of treatment-resistant severe OCD^16,30,36^. Severity of OCD was assessed both pre- and postoperatively using the Yale-Brown Obsessive-Compulsive Scale (Y-BOCS). Postoperative assessment took place 12 months after surgery in Cologne, Grenoble and London cohorts. In case of the London cohort, this followed a four-step clinical trial (2×3 months blinded stimulation at one target followed by 6 months of stimulation at both targets, the last three months using clinically optimized parameters. For details see^16^). In the Madrid cohort, each of the four contact pairs were activated for three months, with a one month wash-out period between trials and a three month sham period. In our analysis, this leads to 32 data points (i.e. stimulation-based outcomes). Patients’ demographic details are provided in Table 1. All patients gave written informed consent. The protocols were approved by the Ethics Committee of each center, respectively.

For all patients in the four cohorts, high-resolution structural T1-weighted images were acquired on a 3.0-Tesla MRI-scanner, before surgery. Postoperative computer tomography (CT) was obtained in thirty-three patients after surgery to verify correct electrode placement, while eleven patients from the Grenoble cohort and the six London patients received postoperative MRI instead. Postoperative MRI parameters were as follows. Grenoble cohort: T1-weighted 3D-FFE scans were acquired on a 1.5T Philips MRI scanner with a 1.0×1.0×1.5 mm^3^ voxel size; TR: 20 ms, TE: 4.6 ms, flip angle: 30 deg. London cohort: T1-weighted 3D-MPRAGE scans were acquired on a 1.5T Siemens Espree interventional MRI scanner with a 1.5×1.5×1.5 mm^3^ voxel size and three-dimensional distortion corrected using the scanner’s built-in module; TR: 1410 ms, TE: 1.95 ms, FOV: 282 mm, flip angle: 10 deg, acquisition time 4 min and 32 s, relative SNR: 1.0.

### DBS Lead Localization and VTA Estimation

DBS electrodes were localized using Lead-DBS software (http://www.lead-dbs.org) as described in^86^ and^83^. Briefly, postoperative CT and MRI scans were linearly coregistered to preoperative T1 images using Advanced Normalization Tools (ANTs, http://stnava.github.io/ANTs/)^88^ Subcortical refinement was applied (as a module in Lead-DBS) to correct for brain shift that may have occurred during surgery. Images were then normalized into ICBM 2009b Nonlinear Asymmetric (“MNI”) template^42^ space using the SyN approach implemented in ANTs^89^, with an additional subcortical refinement stage to attain a most precise subcortical alignment between patient and template space (“Effective Low Variance” preset as implemented in Lead-DBS). This specific method was top performer for subcortical image registrations in a recent comparative study that involved >10,000 nonlinear warps and a variety of normalization techniques^85^. Both coregistrations and normalizations were visually reviewed and refined if needed. DBS electrodes were then localized using Lead-DBS and warped into MNI space.

In the Grenoble, Cologne and Madrid groups, Volumes of Tissue Activated (VTA) were estimated using a finite element method (FEM) as described in^83^. Briefly, a volume conductor model was constructed based on a four-compartment mesh that included gray matter, white matter, electrode contacts and insulated parts. Gray matter was defined by the CIT-168^40^ and DISTAL^41^ atlases for the ALIC-/NAcc and STN-cohorts, respectively. These atlases were specifically adapted or created for use within the Lead-DBS pipeline. The electric field (E-field) distribution was then simulated using an adaptation of the FieldTrip-SimBio pipeline^90^ that was integrated into Lead-DBS (https://www.mrt.uni-jena.de/simbio/; http://fieldtriptoolbox.org/) and thresholded at a level of 0.2 V/m^83^.

For the London test cohort, we chose to use the original VTAs of the published study by Tyagi et al.^16^. These had instead been processed using Medtronic SureTune^™^ software and transferred into MNI space within the original study. The reason we chose to use the original VTAs were twofold. First, it would demonstrate generalizability of our findings (i.e. that our results could still be useful in case electrodes were localized using different software). Second, we aimed at yielding maximal transferability to the study by Tyagi et al. which reported on the rich London dataset in more depth.

### Connectivity Analysis

Structural connectivity between VTAs and all other brain areas was calculated based on a normative connectome as similarly done in previous work^29,30,41,44,80,83^. Specifically, a whole-brain connectome based on state-of-the-art multi-shell diffusion-weighted imaging data from 985 subjects of the Human Connectome Project (HCP) 1200 subjects data release^81^ was calculated in each patient using Lead-Connectome. Whole-brain fiber tracts were then normalized into standard space using a multispectral warp based on T1-weighted, T2-weighted, and diffusion-weighted acquisitions using ANTs (using the same “Effective Low Variance” preset implemented in Lead-DBS). In each subject, a total of 6,000 fibers were sampled and aggregated to a joint dataset in standard space, resulting in a set of 6,000,000 fibers across 985 HCP subjects. For each of these tracts, a “Fiber T-score” was assigned by associating the fiber tract’s connectivity to VTAs across patients with clinical outcome (Figure 7). Specifically, (mass-univariate) two-sample t-tests between clinical outcomes in connected and unconnected VTAs were performed for all 6,000,000 tracts. Needless to say, these T-scores were not meant to result in significant results but instead formed a model that could be used for out-of-sample predictions in other DBS cohorts. T-values from these tests and could be positive or negative (since two-sided t-tests were performed). A high absolute T-value meant that the fiber was strongly discriminative or predictive for clinical outcome. For instance, a tract that was connected exclusively to VTAs in good responders (and not to VTAs of poor responders) would receive a high positive score. In return, a patient would most likely show more pronounced clinical benefit, if her/his VTA was strongly connected to many fibers with high positive T-values but not to many with negative scores. This analysis made it possible to assign aggregated “Fiber T-scores” to each VTA in subsequent prediction analyses.

**Figure 7.**
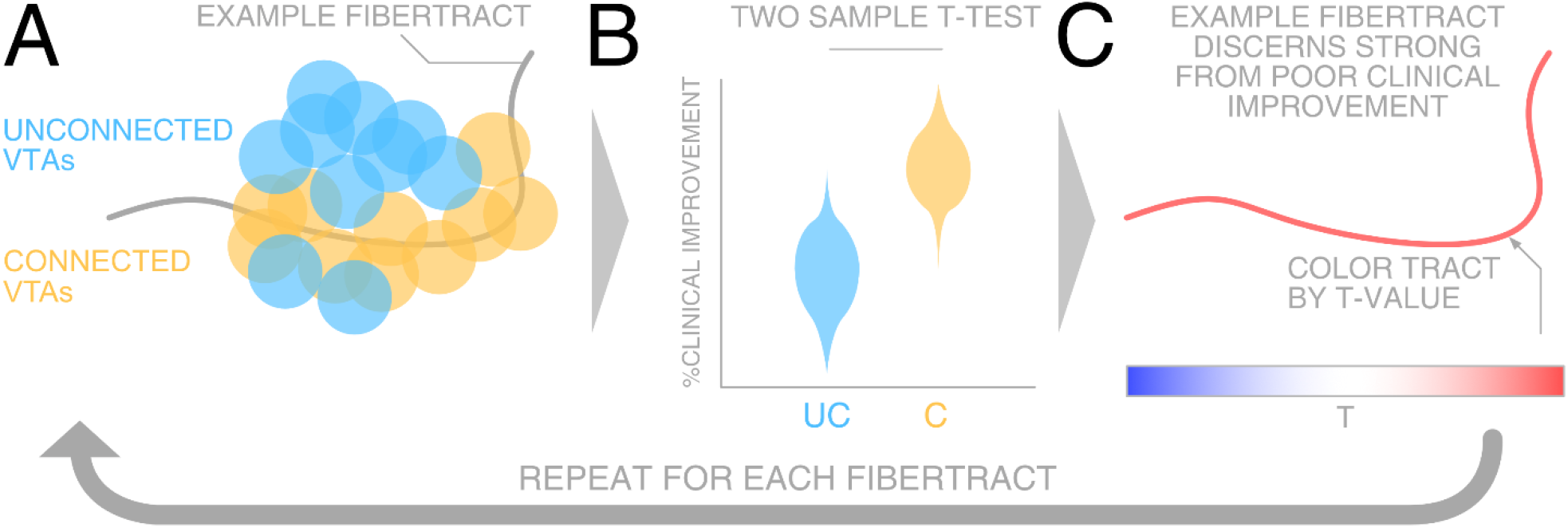
Summary of methods to define a weight for each tract. A) For each fiber, VTAs were grouped into either connected (C; yellow) or unconnected (UC; blue) sets across patients. B) Two-sample t-tests between clinical improvements in connected and unconnected VTAs were calculated in a mass-univariate fashion for each fiber tract separately. C) The resulting T-value of this analysis leads to the “weight” that each fiber was given, as well as the color in visualizations throughout the manuscript. Here, red means that the fiber tract is favorably connected to good responders while blue indicates the opposite (and the saturation of tracts denotes how discriminative they are).

To account for the fact that larger VTAs would potentially automatically receive higher Fiber T-scores, these were divided by the stimulation amplitude throughout the manuscript. Finally, Monte-Carlo random permutations (× 1000) were conducted to obtain *p*-values, except for two-sample t-tests. This procedure is free from assumptions about the distributions (e.g. Student-T for R-values) which are typically violated in small sample sizes^91^. Scatterplots are visualized with 95% confidence bounds (gray or light-red areas).

## Data and code availability

The DBS MRI datasets generated during and analyzed during the current study are not publicly available due to data privacy regulations of patient data but are available from the corresponding author on reasonable request. The resulting tract atlas and code used to analyze the dataset is openly available within Lead-DBS /-Connectome software (https://github.com/leaddbs/leaddbs).

## Acknowledgements

We would like to thank Kristy Kultas-Ilinsky and Igor Ilinsky, Rudolf Nieuwenhuys and Suzanne Haber for counseling regarding the anatomical validity of tractography results presented in this study. We would like to thank Cyril Pernet, Wolf-Julian Neumann, Roxanne Lofredi for consultation and help on the manuscript.

This work was supported by the German Research Foundation (Deutsche Forschungsgemeinschaft, Emmy Noether Stipend 410169619 to AH, SPP 0141 to AAK and KFO 219 to JK). Data were provided in part by the Human Connectome Project, WU-Minn Consortium (Principal Investigators: David Van Essen and Kamil Ugurbil; 1U54MH091657) funded by the 16 NIH Institutes and Centers that support the NIH Blueprint for Neuroscience Research; and by the McDonnell Center for Systems Neuroscience at Washington University.

## Author contributions

N.L. and A.H. conceptualized the study. N.L. and A.H. developed the software pipeline used, analyzed data and wrote the manuscript. J.C.B. conceptualized the study, acquired patient data and revised the manuscript. S.T. performed literature analyses and wrote the manuscript. H.A. acquired and processed patient data and revised the manuscript. G.J.B.E., A.B., and A.M.L. processed and analyzed human connectome data and revised the manuscript. B.A. processed patient data, conceptualized part of the study and revised the manuscript. A.K., B.S., J.B., L.Z., E.J., S.C., V.V.V., M.P., J.K. and A.A.K. acquired patient data and revised the manuscript.

## Materials & Correspondences

Correspondence and material requests should be addressed to Ningfei Li

## Supplementary Material

**Figure S1.**
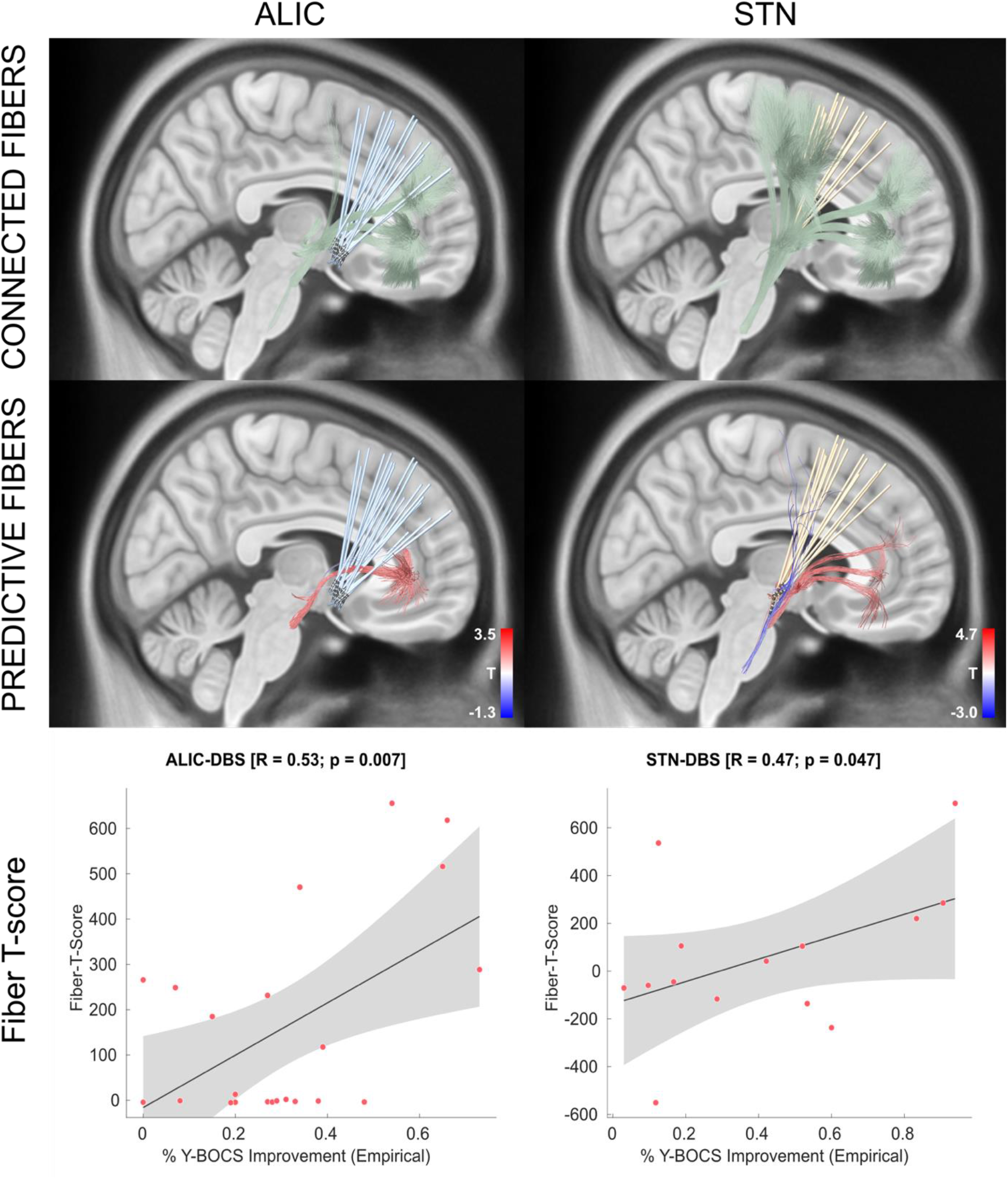
Replication of Figure 2 using the basal ganglia pathway atlas^22^ instead of a normative connectome. This atlas is not based on diffusion-weighted imaging but on established anatomical expert knowledge and thus free of false-positive connections. While the STN target isolates three distinct bundles that traverse within the ALIC, the only predictive bundle isolated by the ALIC electrodes is the hyperdirect pathway connecting dACC and STN. In this context, it is crucial to note that connections from other regions of the prefrontal cortex (such as the ventrolateral or medial prefrontal cortex) to the STN were not represented in the atlas (and could thus represent false-negative findings, as shown by the white-matter gaps between atlas components).

**Figure S2.**
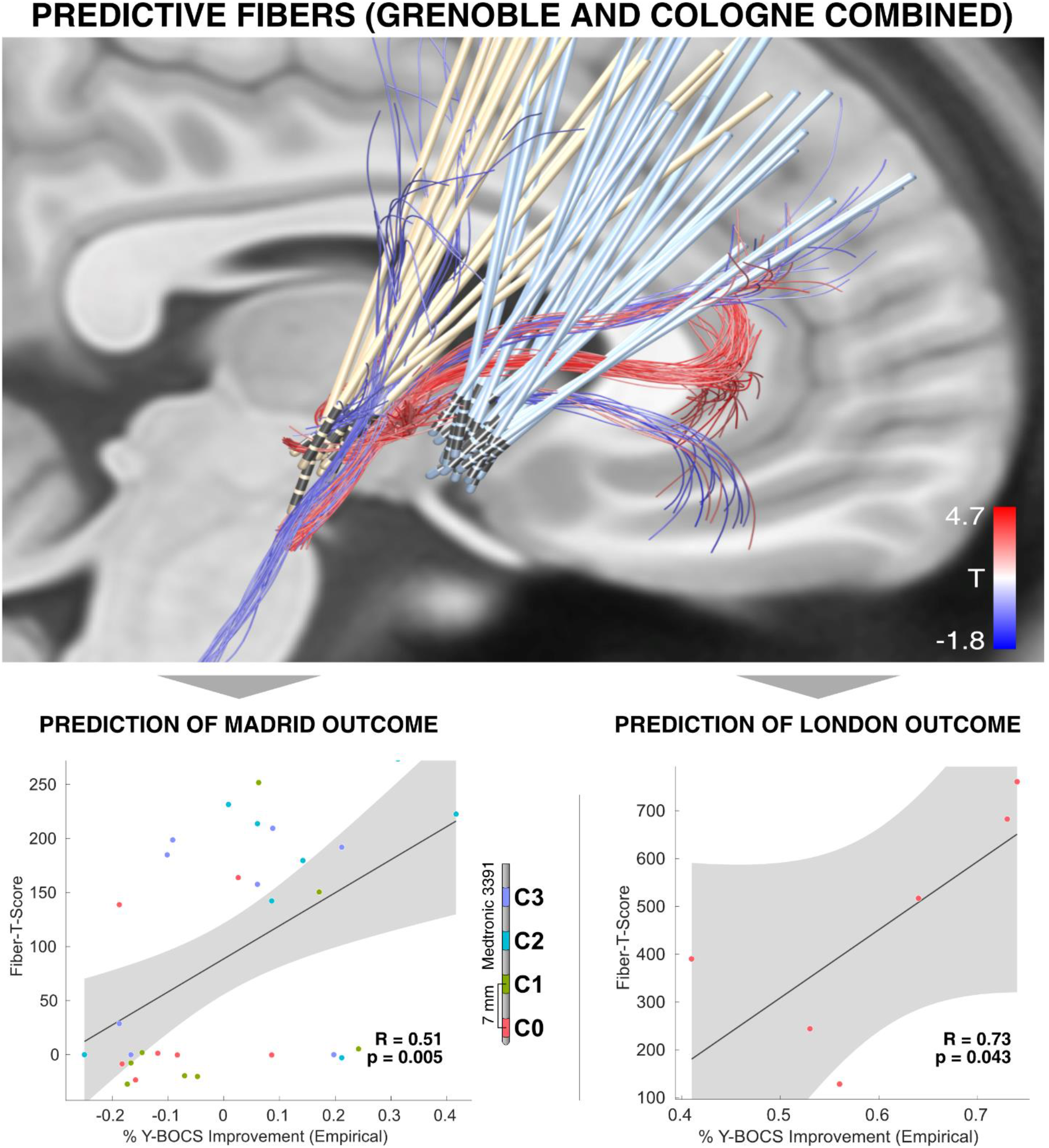
Replication of Figure 4 using the basal ganglia pathway atlas^22^ instead of a normative connectome. This atlas is not based on diffusion-weighted imaging but on established anatomical expert knowledge and is thus free from false-positive connections. The most predictive bundle present in the atlas was the hyperdirect pathway connecting dACC and STN. In this context, it is crucial to note that connections from other regions of the prefrontal cortex (such as the ventrolateral or medial prefrontal cortex) to the STN were not represented in the atlas (and could thus represent false-negative findings, as shown by the white-matter gaps between atlas components).

**Figure S3.**
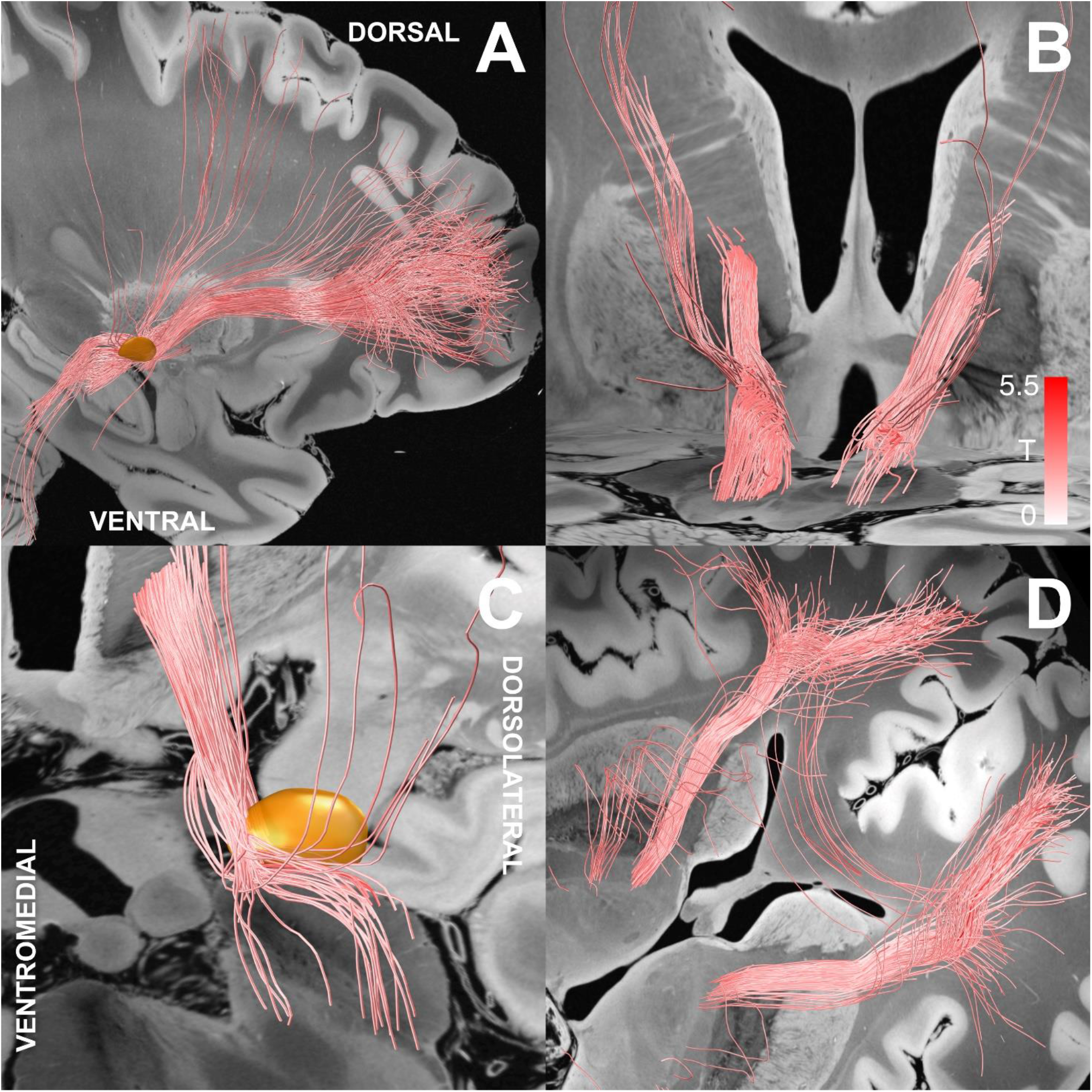
Positively predictive fiber tracts that were discriminative in both ALIC- and STN-cohorts shown from multiple angles to further characterize them, anatomically. A) Sagittal overview with STN in orange. B) Trajectory within the internal capsule while it passes putamen and caudate. C) Close-up showing the tracts course within and below the anterior STN. D) Oblique view of the axial aspect after entering the anterior limb of the internal capsule coming in from ventrally. A 7T 100 um postmortem template is shown in the background^92^.

**Figure S4.**
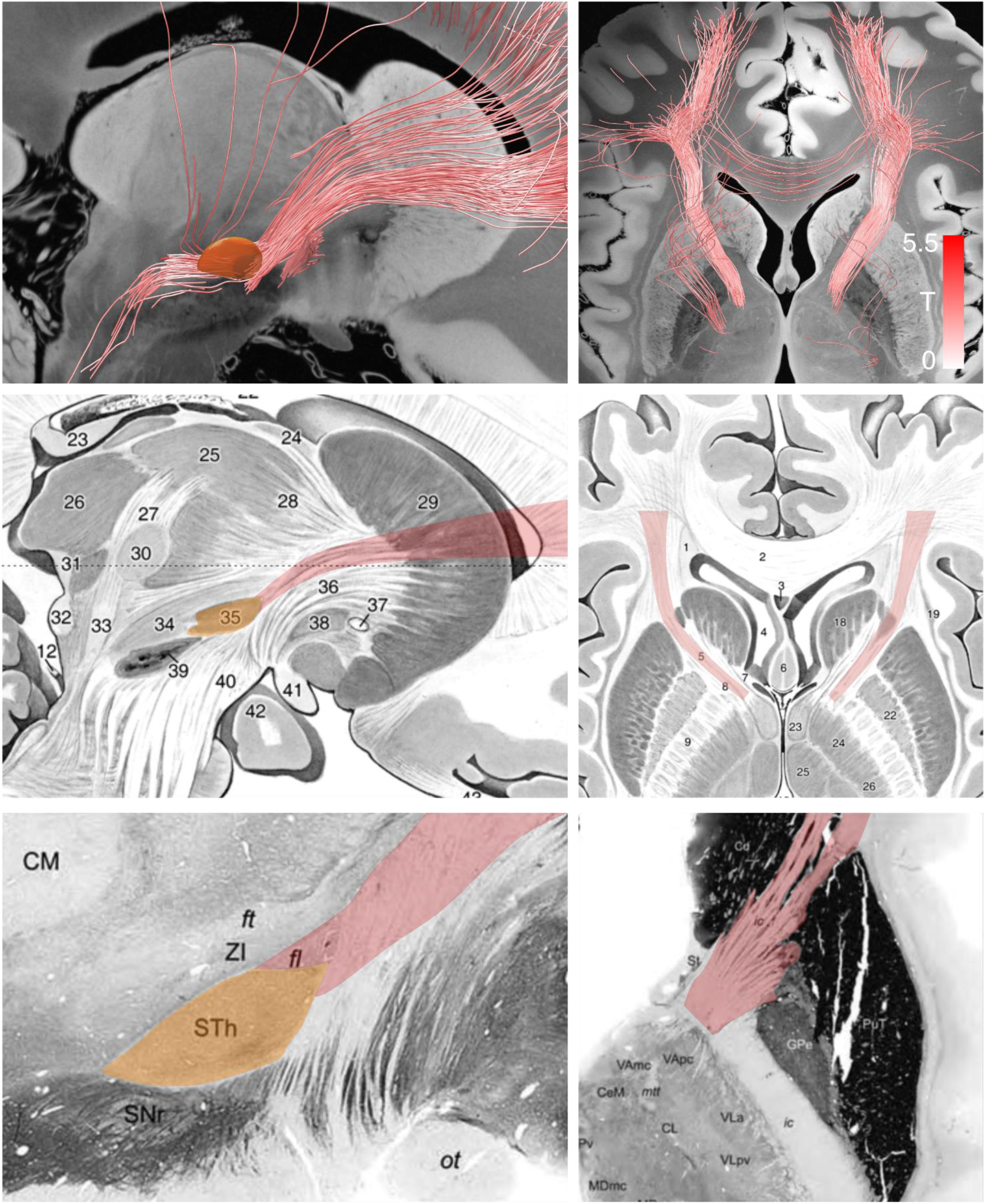
Direct synopsis of the tract target (top row) with anatomical white-matter textbook definitions. Backdrop in top row shows a 7T 100 um postmortem template^92^. Mid row shows data adapted from^46^, Bottom row from^93^.

### Literature based DBS target analysis (cf. Table 2, Figure 5)

For each publication (Table 2), pre- and postoperative average Y-BOCS scores were extracted to calculate the average percent change in Y-BOCS (difference between avg. preop- and postop-scores divided by avg. preop scores). As in all “meta-analysis” situations, due to differences in study design and reported data, we had to decide which exact values to use in some cases (below).

In 4 studies (Mallet et al. 2008, Tsai et al. 2010, Nuttin et al. 2003, Maarouf et al. 2016), improvements were unambiguously reported. Tsai et al. reported one case of hypomania induced by ventral capsule stimulation (in the vicinity of NAcc). Thus, we used the reported postoperative Y-BOCS score collected about one month after surgery to ensure consistency of pre- and postoperative medication status (sodium valproate dose was later changed to ameliorate hypomanic side effects induced by higher stimulation amplitudes). Nuttin et al. reported average postoperative Y-BOCS scores that had been maintained for 21 months (Nuttin et al. 2003). In one study (Maarouf et al. 2016) that included four cases operated with different targets, we used average postoperative Y-BOCS scores of the latest follow-up in order to calculate the average percentage change in Y-BOCS. These applied to the MD/VA group of active contacts. Still, MD coordinates were used in the correlation analysis since all patients had most of their contacts in the MD region (and MD and VA nuclei are close to each other). Besides, follow-up Y-BOCS scores were not labelled as belonging to a specific target (MD vs. VA) based on the information reported. For two studies (Coenen et al. 2017 and Lee et al. 2019), percentage change in Y-BOCS scores were directly reported. Coenen et al. reported two OCD patients implanted to the sl-MFB target (Coenen et al. 2017). While one patient had an improvement of 50%, the other was reported to have *over* 35% improvement. Thus, we averaged 50 and 35 % to obtain a publication based average percent change in Y-BOCS. Lee et al. reported two average Y-BOCS improvements (52% for 1-year follow-up and 54% for latest follow up; Lee et al. 2019). We used the 1-year follow up improvement in Y-BOCS to be consistent with the other studies.

This led to average literature-based improvements in six publications for correlation analysis, while three studies (Nair et al. 2014, Strum et al. 2003 and Nuttin et al. 2013) did not report Y-BOCS improvement scores. These six average improvement values were correlated with weighted overlaps between the reported average stimulation sites and the tract-target identified here. To do so, stereotactic coordinates were converted to MNI space using a novel probabilistic method^44^. A sphere of radius 3 mm was introduced at this site and heavily smoothed with a sigma of 6 mm (to allow for a weighted/distance measure with the tract). Weighted overlap values between these smoothed volumes and the T-values of the tract-target were multiplied to derive a literature-based Fiber-T-score which was correlated with average improvement scores.

